# Growth rate regulates RNA polymerase II transcription elongation in *Saccharomyces cerevisiae*

**DOI:** 10.1101/2024.11.26.625403

**Authors:** A.I. Garrido-Godino, R. González, M. Martín-Expósito, S. Chávez, J.E. Pérez-Ortín, F. Navarro

## Abstract

Cells must adapt to changing environmental conditions to maintain their fitness and to compete with other genotypes during the natural selection process. The growth rate (GR) is a determining factor in this competition, and it influences gene expression. Some genes increase mRNA levels, while others decrease with the GR. mRNA levels depend on the dynamic balance between their synthesis by RNA polymerase II and their degradation rates. RNA polymerase I and III are also influenced by the GR because they transcribe protein synthesis machinery required to make proteins that increase cell mass during growth. Although RNA levels have been extensively studied in relation to the GR in many organisms, synthesis and degradation rates have, however, been much less investigated. In a previous work, we found a positive correlation between RNA polymerase (RNA pol) II transcription and mRNA degradation with GRs in yeast in batch cultures. Here we extend our study under constant growth conditions in a chemostat and find that overall chromatin-associated RNA pol II levels increase in parallel with the GR. This increase appears to involve the accumulation of partially dephosphorylated RNA pol II with a greater tendency to backtracking, which suggests that the GR regulates RNA pol II-dependent transcription at the elongation level. RNA pol I also increases its association with chromatin with the GR, which confirms the general dependence of at least RNA pol I and II transcription on the GR.

## INTRODUCTION

Growth is an inherent property of living things [1]. As growth implies not only an increase in both volume and mass, cells must synthesise new molecules to cope with this need. Given that proteins constitute a large part of the cell mass [2, 3], most of this effort is spent on protein synthesis (revised in [2]). The rate at which cells grow should, therefore, be closely linked with protein synthesis. However, as proteins are translated from a template molecule, mRNAs level and the ribosome/mRNAs ratio seem to be critical for correct protein synthesis [2], and the cellular growth rate (GR) should play a pivotal role in regulating gene transcription (discussed in [2, 4–7]). This relation is multifaceted and dynamic, and it impacts various cellular function and gene expression aspects. The GR affects all three eukaryotic RNA polymerases (RNA pol). A faster GR often correlates with increased transcriptional activity due to heightened metabolic demands and the need for additional cellular components. At higher GRs, cells typically require more mRNA for protein synthesis, which leads to a need for high RNA pol II transcription levels. To translate these additional mRNAs, a corresponding increase in translational machinery is necessary that, in turn, requires larger amounts of rRNA and tRNAs.

To increase or decrease mRNA levels, cells can use either transcription or mRNA decay machineries [8]. Finally, increased mRNA levels at higher GRs encompasses most genes, but it is especially relevant for those genes associated with cell division and biosynthesis pathways [5–7]. Conversely, slower GRs may prompt cells to down-regulate global mRNA, rRNA and tRNA levels (and concomitantly their synthesis rates: SR) to preserve energy and resources. Down-regulation in GRs can trigger signalling pathways and cellular stress responses that, in turn, affect transcriptional regulation. For instance, nutrient availability, environmental cues or stress conditions can modify GRs and, consequently, modulate transcriptional programmes to adapt to changing circumstances [4, 7]. Although mRNA levels have been extensively studied in relation to GRs in many organisms [5–7], synthesis and decay rates have, however, been much less investigated. We previously used budding yeast *Saccharomyces cerevisiae* to study gene-specific and global changes in RNA pol II synthesis and decay rates [4, 8, 9]. We found that, as expected, the global SR increases with the GR. On the contrary and surprisingly, although levels of some specific mRNAs increase or decrease their levels, global mRNA levels were mostly invariant due to a compensatory global increase in mRNA decay rates [4]. The experimental setup in those studies used batch cultures of different yeast strains growing at specific GRs in the exponential growth phase. Employing batch cultures limits the possibility of having well-defined and controlled GRs. Moreover, the use of mutant strains with different physiological defects adds some noise to the observed results. Nevertheless, cell physiology can be studied in a more controlled manner with continuous cultures, such as chemostats, which allow constant GRs to be maintained [10–12]. Furthermore, accurate GR regulation enables their alignment between wild-type (WT) and mutant cultures with slower GRs or among various strain backgrounds. This is achieved by maintaining cells in a constant growth state that is below the maximum GR for both cultures [12]. Therefore, we decided to use chemostat cultures of WT yeasts to check the dependence of RNA pol II global transcription on culture GRs. We also applied new approaches to evaluate the amount and concentration of RNA pol II in chromatin as a proxy to the RNA pol II SR. We found that global RNA pol II chromatin-associated levels increase in parallel to the GR, which confirmed our previous results [4]. This increase appears to involve the accumulation of partially dephosphorylated RNA pol II with a greater tendency to backtracking, which suggests that the GR regulates RNA pol II-dependent transcription at the elongation level.

## RESULTS

### Experimental Design

In order to obtain yeast cells that grow at different GRs, under otherwise similar growth conditions a series of chemostat cultures of strain LMY5.2 [13] was established. Dilution rates (equalling the target GR) ranged from 0.26 to 0.49. The samples for chromatin analyses were taken in the steady state as described in Materials and Methods. The residual glucose values in the steady state ranged from 0.83 g/L to 1.59 g/L, ethanol from 0.91% to 0.17% (v/v), and biomass from 4.06 to 1.01 OD_600_ (Figure 1).

**Figure 1.**
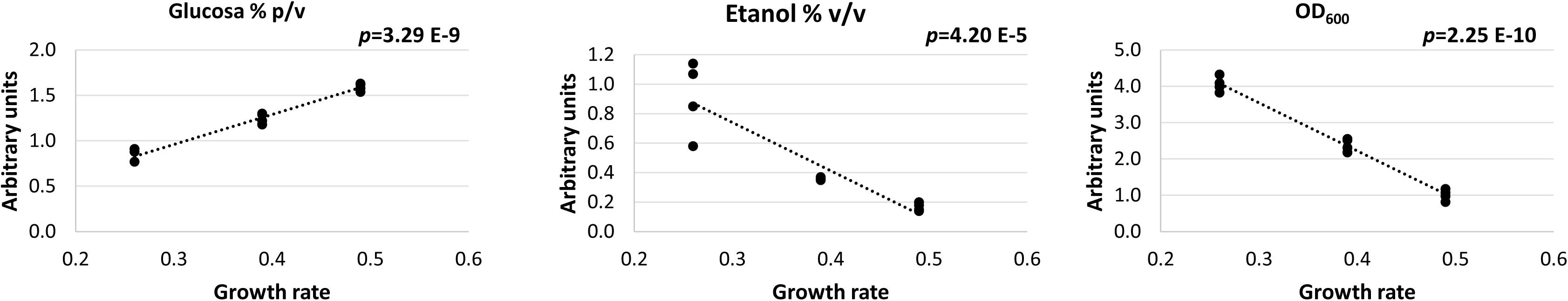
Different parameters corresponding to the yeast cells growing at different growth rates. The data for residual glucose (%, w/v), ethanol (%, v/v) and biomass at OD_600_, from a series of chemostat cultures of strain LMY5.2 grown at several dilution rates (equalling the target GR) ranged from 0.26 to 0.49 (0.26, 0.39 and 0.49).

We wondered whether cell volume correlated with the GR. To find out, by flow cytometry we analysed the forward scatter (FSC) because its intensity is proportional to the cell’s diameter. Accordingly, FSC-A median parameter was used to correlate with cell size:. The analysis of the cells from three different chemostat cultures at distinct GRs from 0.26 to 0.49 h^-1^ (Figure S1) showed that cell size decreased with the GR under our experimental conditions, which agrees with our previous results [4].

### Chromatin-associated RNA pol II increases with the growth rate

In order to investigate whether the amount of RNA pol II varied with the GR, we determined the chromatin-associated RNA pol II levels in the chromatin-enriched fractions prepared by the yChEFs procedure [14, 15] from the same previously employed chemostat cultures at distinct GRs from 0.26 to 0.49 h^-1^. RNA pol II was analysed by western blot with different antibodies against several RNA pol II subunits or epitopes: the anti-Rpb1 (using two different antibodies, the first against the carboxy-terminal domain, CTD, and the second one against the N-terminal domain), anti-Rpb3 and anti-Rpb4 antibodies. A histone H3 antibody was used as a control of the chromatin fraction and a phosphoglycerate kinase (Pgk1) antibody as a control of the cytoplasmic proteins.

As cell size decreased with the GR (see Figure S1), we represented the relative amount of each chromatin-associated RNA pol II subunit in relation to histone H3. This allows to normalise the amount per cell given that the histone content of each cell is fixed and does not depend on cell size [16]. Then we divided it by cell size at each GR (to, thus, represent its cellular concentration) to gain a proxy of the actual SR (see [17] for further explanations) of RNA pol II (Figure 2). The results corroborated the increase in chromatin-bound RNA pol II concentration (by representing the global amount of molecules engaged in transcription) with the GR. This finding suggests an increase in the SR with the GR, as previously reported [4]. The rise in the SR was not proportional to that in the GR, i.e. it is on a sublinear scale (subscaling).

**Figure 2.**
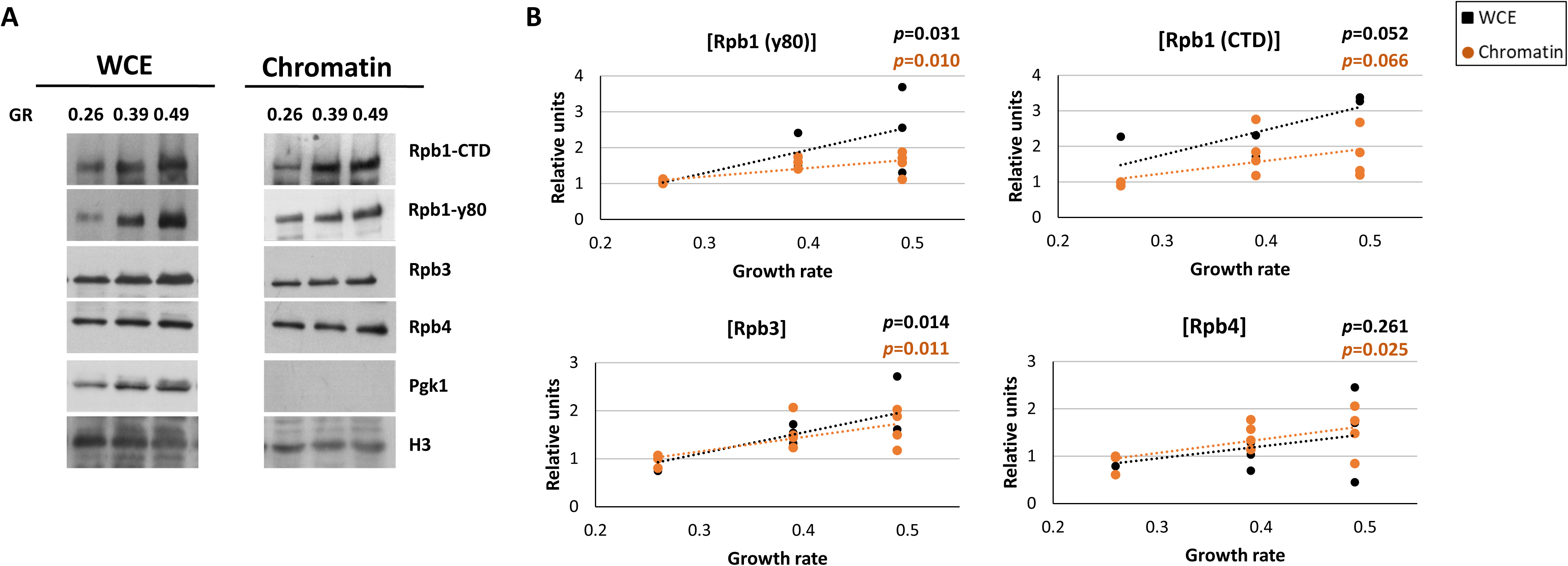
RNA pol II levels in chemostat cultures at different growth rates. **A)** The RNA pol II levels from the chemostat cultures at different growth rates (GRs) were determined by western blot analysis of subunits Rpb1, Rpb3 and Rpb4 in the chromatin-enriched fractions and whole-cell crude extracts (WCEs) isolated by the yChEFs procedure [14, 15]. Two different anti-Rpb1 antibodies (one against CTD and another, y80, against the terminal domain), anti-Rpb3 and anti-Rpb4 antibodies were used. Histone H3 and Pgk1 were employed as the internal controls of chromatin and cytoplasm, respectively. **B**) The relative amounts of Rpb1 (y80 antibody), Rpb3 and Rpb4 subunits vs. Histone H3 (shown in A) were divided by cell volume (see Figure S1) to represent RNA pol II subunit as cellular concentrations. A Multiple Regression model was utilised for the statistical analysis and the associated p-values are shown (statistical significance, p-value < 0.1).

We also analysed the Rpb1, Rpb3 and Rpb4 concentrations (normalised to Histone H3 and dividing it by cell size at each GR) in the corresponding whole-cell crude extracts (WCEs). Differences with the GR were noted, which were statistically significant, and with the only exception of Rpb4 (Figure 2). These data indicate that the amount of RNA pol II engaged in transcription increased with the GR, probably caused by the increase in its cellular concentration.

We further analysed the concentrations of RNA pol I and III because they transcribe rRNAs and tRNAs, the most abundant RNAs in cells necessary for protein synthesis, and also because their SRs have been previously shown to correlate with cellular GRs [4, 18]. To do so, by western blot we analysed the levels of chromatin-associated RNA pol I and III in the above-employed samples with antibodies against the Rpa34 and Rpc53 subunits of RNA pol I and III, respectively. As shown in Figure 3, and as expected, the chromatin-associated RNA pol I concentration positively correlated with the GR. In this case, the increase in the SR with the GR was faster than for RNA pol II, and came closer to being linear. However, no significant differences in the Rpc53 subunit concentration in WCEs or the chromatin fractions were observed along the GR (Figure 3).

**Figure 3.**
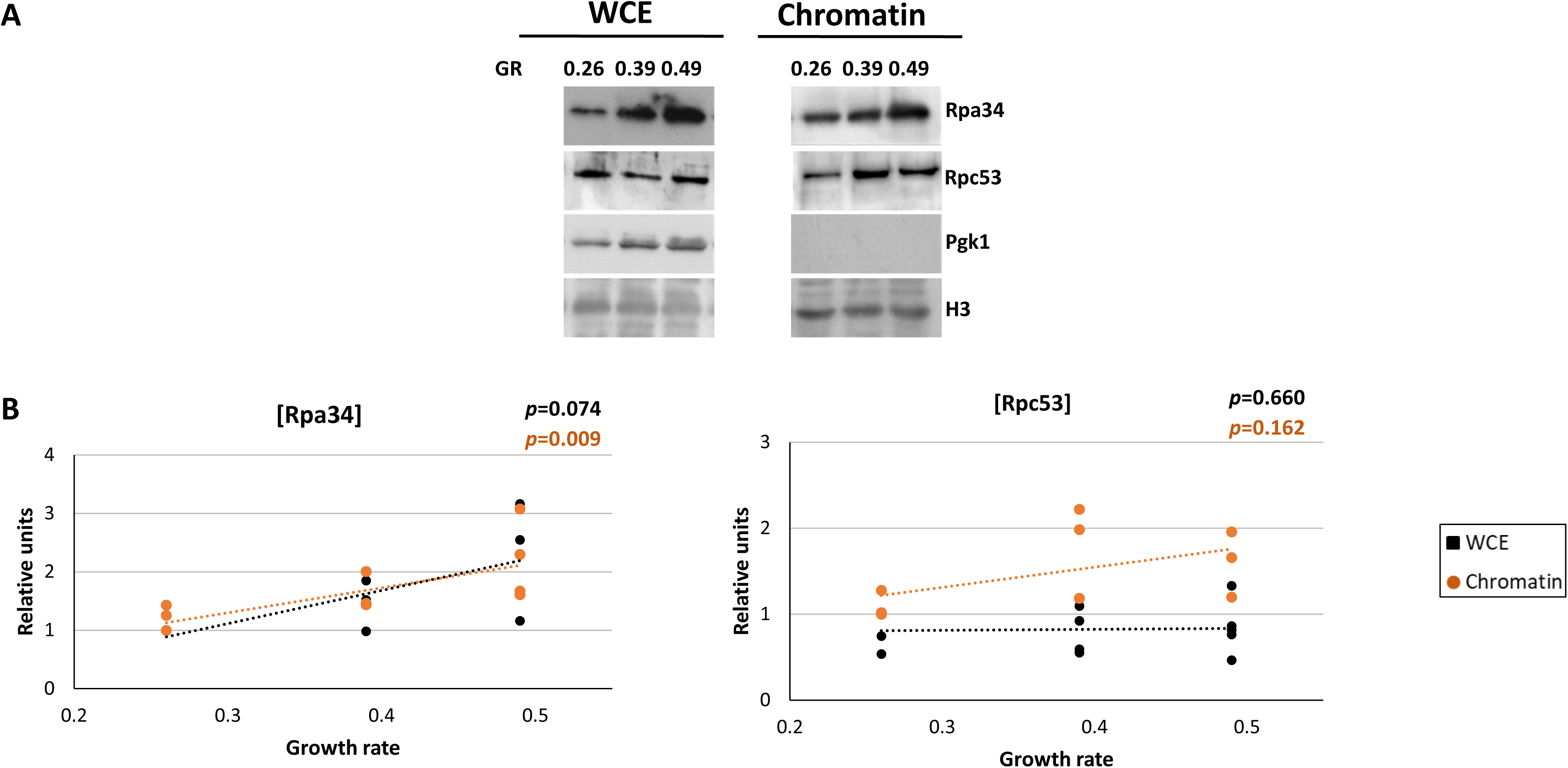
RNA pol I and III levels in chemostat cultures at different growth rates. **A)** The RNA pol I and III levels from chemostat cultures at different growth rates (GRs) were determined by western blot analysis of the Rpa34 (RNA pol I) and Rpc53 (RNA pol III) subunits in the chromatin-enriched fractions and whole-cell crude extracts (WCEs) isolated by the yChEFs procedure [14, 15]. Anti-Rpa34 and anti-Rpc53 antibodies were used. Histone H3 and Pgk1 were employed as internal controls of chromatin and cytoplasm, respectively. **B)** The quantification of the data in A for each RNA pol I and III subunits normalised against histone H3 is represented as cellular concentrations (see Figure 2 legend). Four biological replicates were used for each GR. Data represent median and standard deviation. A Multiple Regression model was used for the statistical analysis and the associated p-values are shown (statistical significance, p-value < 0.1).

### Phosphorylation of elongating RNA pol II molecules decreases with the growth rate

Elongating RNA pol II is characterised by the phosphorylation of its CTD domain. We investigated if an increased GR would bring about any change in this phosphorylation. To do so, we used the same samples as in the previous analyses to perform the western blot analysis with an antibody that detects the phosphorylated Ser2 residues of Rpb1 CTD (Figure 4), a hallmark of transcription elongation [19]. We also analysed two other posttranslational modifications of the RNA pol II CTD: Ser5-P and Ser7-P. These two phosphorylation marks are established during transcription initiation and, while Ser5-P levels lower during elongation, Ser7-P tends to be sustained throughout the elongation process [20, 21] As shown in Figure 5B, the Ser2-P and Ser5P concentrations did not increase with the GR; on the contrary, the Ser7-P concentration drastically dropped. We then represented the Ser2P/Ser7P to Ser2P/Ser5P ratio (Figure 4C). As shown, the Ser2P/Ser7P and Ser2P/Ser5P ratios evidenced a clear increase with the GR, which suggests a strong regulation of CTD phosphorylation in response to GR. However, and importantly, albeit at different rates all the CTD-phosphorylated forms of RNA pol II decreased with regards to the total amount of the total chromatin-associated RNA pol II levels (Figure 4D).

**Figure 4.**
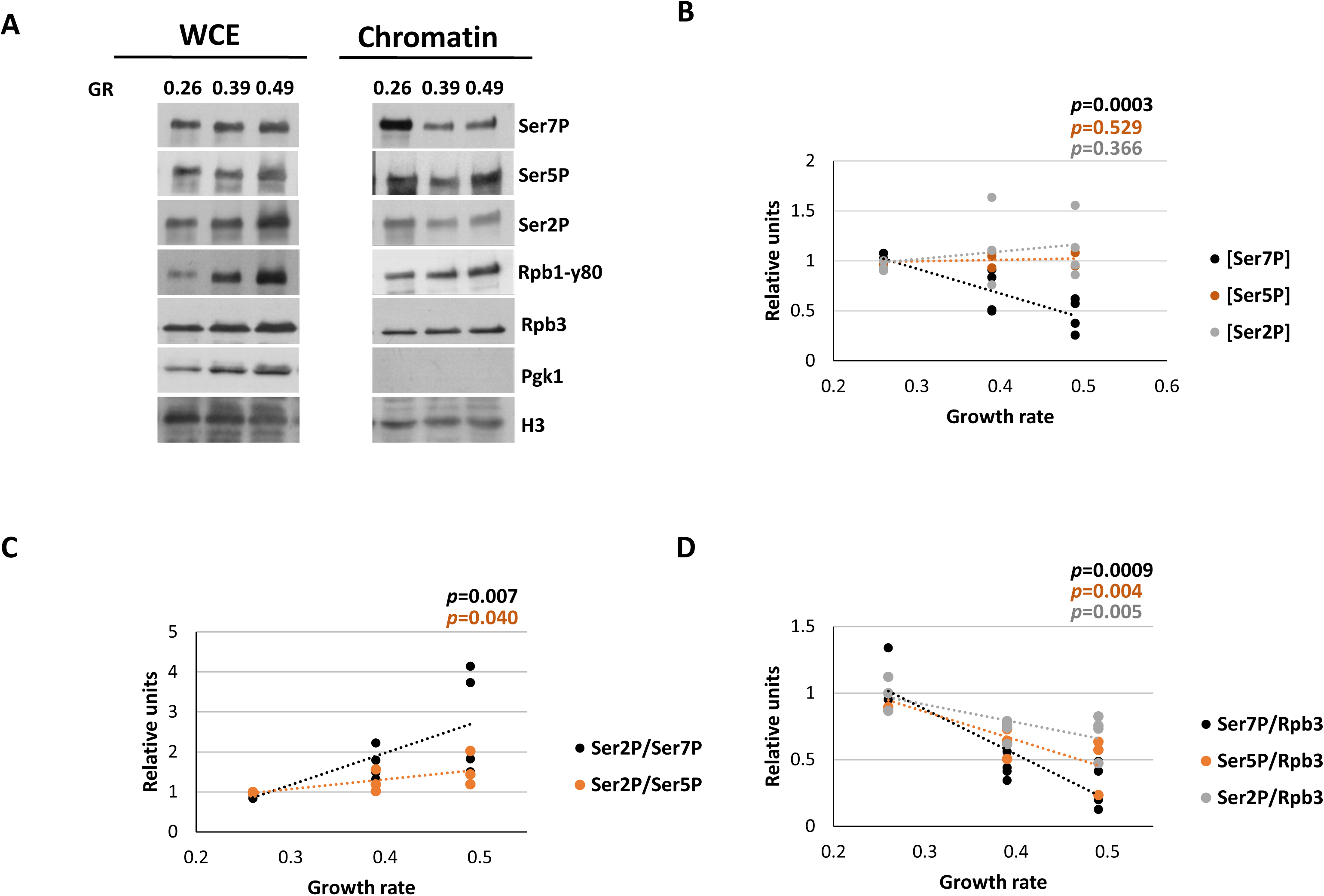
RNA pol II CTD phosphorylation levels in chemostat cultures at different growth rates. **A)** CTD-Ser2, -Ser5 and -Ser7 phosphorylations, as well as the Rpb1 levels from the chemostat cultures at different growth rates (GRs), were determined by western blot analysis in the chromatin-enriched fractions and whole-cell crude extracts (WCEs) isolated by the yChEFs procedure [14, 15] with specific antibodies. Histone H3 and Pgk1 were used as internal controls of chromatin and cytoplasm, respectively (see Figure 2 legend). **B)** Ser2P-CTD (representative of elongating RNA pol II), Ser5P-CTD and Ser7P-CTD normalised over Histone H3 (divided by cell volume (see Figure S1)) from the chromatin fractions to represent the Ser2P-, Ser5P- and Ser7P-CTD concentration, corresponding to the westerns appear in A. **C)** Quantification ratios from the westerns blot for Ser2-CTD/Ser7-CTD and Ser2-CTD/Ser5-CTD over cell size (FSC-A median). **D)** Quantification ratios from the westerns for Ser2P-CTD/Rpb3, Ser5P-CTD/Rpb3 and Ser7P-CTD/Rpb3. Four biological replicates were used per GR. Data represent median and standard deviation. A Multiple Regression model was used for the statistical analysis and the associated p-values are shown (statistical significance, p-value < 0.1).

**Figure 5.**
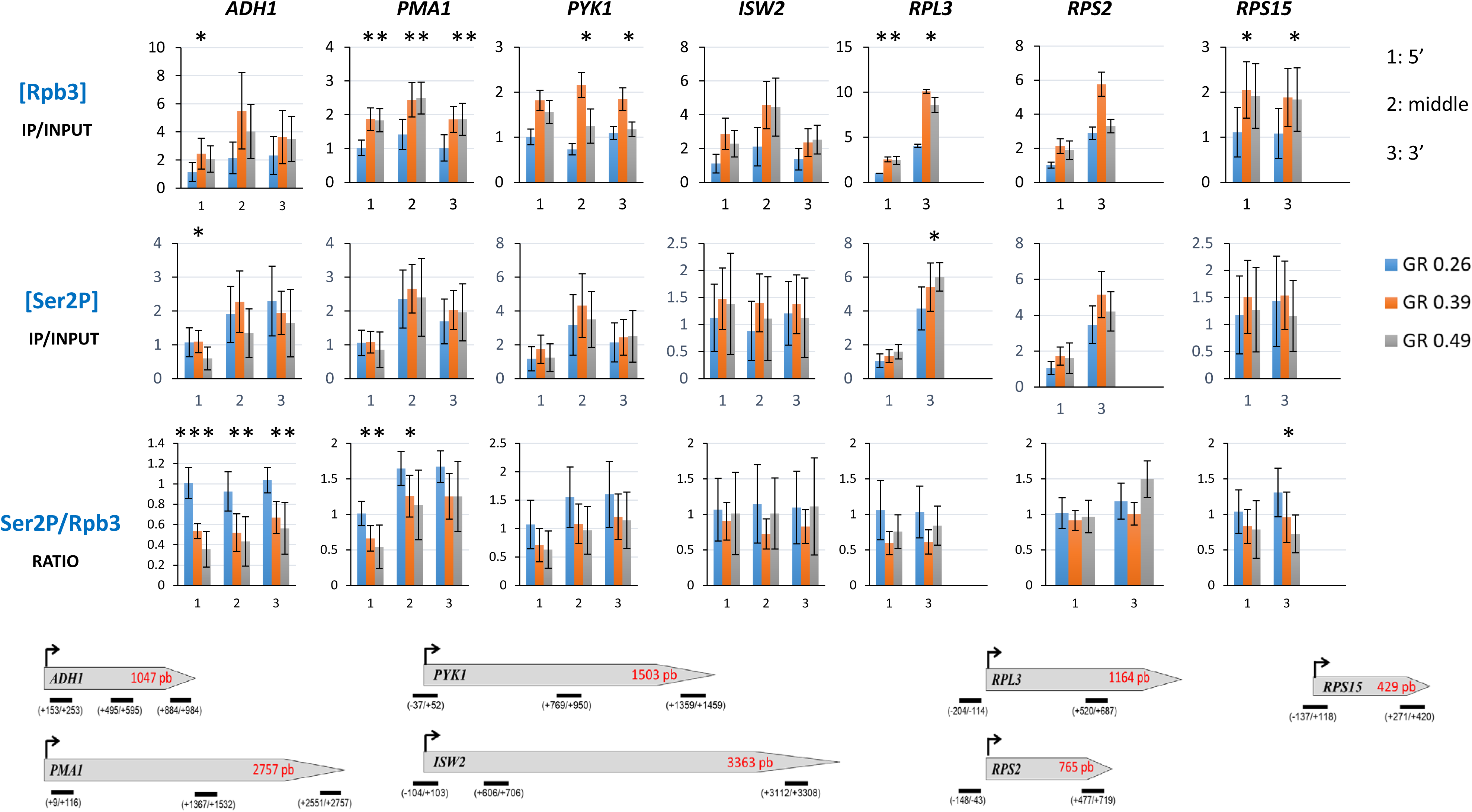
RNA pol II and RNA pol II CTD-Ser2phosphorylation levels in chemostat cultures at different growth rates, analysed by Chromatin Immunoprecipitation. Chromatin immunoprecipitation (ChIPs) analyses for Rpb3 and Ser2P-CTD occupancy in different transcription units from chemostat cultures at different growth rates (GRs). Representation of the ratios between the data for the Ser2P-CTD and Rpb3 analyses are also shown. Data denote IP/INPUT ratios. At least three biological triplicates were used per GR. The position of each analysed amplicon, along each transcription unit, are graphically represented on the bottom. A multiple regression model was used for the statistical analysis and the associated p-values are shown (*: p-value < 0.1; **: p-value < 0.05; ***: p-value < 0.005).

Altogether, these results suggest that the overall GR-dependent increase in RNA pol II present in chromatin involves alterations of this enzyme during elongation, as reflected by its CTD phosphorylation marks.

To investigate whether the data shown above for whole chromatin could be confirmed by individual genes analysis, we decided to study the occupancy of the total RNA pol II in individual genes by chromatin immunoprecipitation (ChIP) using samples from the same cultures as in the previously presented data. We analysed several amplicons along the transcribed region of the different genes, from 5’ to 3’. As Figure 5 depicts, total RNA pol II occupancy increased with the GR for most of the analysed transcription units.

We also determined by ChIP the occupancy of Ser2-P RNA pol II, which represents the amount of Ser2P phosphorylation that corresponds to elongating RNA pol II. As shown in Figure 5, although total RNA pol II occupancy significantly increased with the GR for most the analysed transcription units, the occupancy of Ser2-phosphorylated RNA pol II did not change significantly with the GR. Accordingly, and in line with previous data, the phosphorylation level of elongating RNA pol II (Ser2-P/Rpb3) lowered with the GR in some analysed genes (Figure 5).

All these data support data obtained with chromatin fractions, and confirm that increased GR promote higher transcriptionally-engaged RNA pol II levels with a lower degree of phosphorylation.

### The total concentration of the mRNA degradation factors, but not their association with RNA pol II in chromatin, increases with the growth rate

Global mRNA turnover increases with the GR [9] and decreases with cell volume in budding yeast [22]. This process is regulated by some proteins that participate in both mRNA synthesis and degradation, such as the major deadenylase complex, Ccr4-Not and 5’-3’exonuclease Xrn1 [23–25].

To analyse whether proteins involved in mRNA degradation could change their concentration with the GR, by western blot we analysed the amount of Ccr4 and Xrn1 in both WCEs and chromatin in the same samples used before. Interestingly, we observed an increase in the [Ccr4] and [Xrn1] in the WCEs with the GR, which was slighter for chromatin fractions (Figure 6A and B). Furthermore the Xrn1 to RNA pol II ratio in chromatin seemed to lower with the GR (Figure 6C).

**Figure 6.**
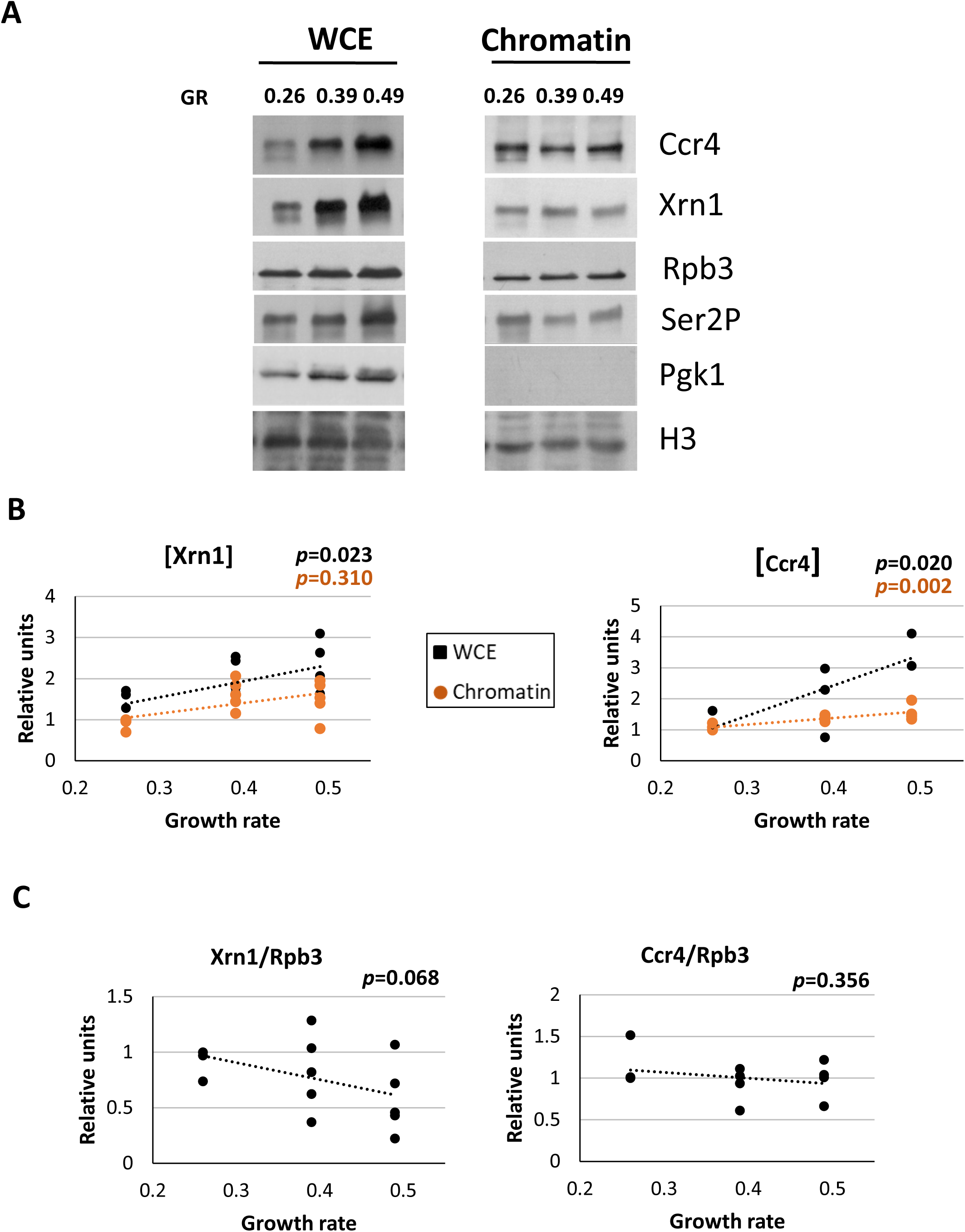
Ccr4 and Xrn1 levels in chemostat cultures at different growth rates. **A**) The Ccr4 and Xrn1 levels from chemostat cultures at different growth rates (GRs) were determined by western blot analysis in the chromatin-enriched fractions and whole-cell crude extracts (WCEs) isolated by the yChEFs procedure [13, 14]. Histone H3 and Pgk1 as internal controls of chromatin and cytoplasm, respectively. **B**) Quantification of the western blot analysis shown in A for Xrn1 and Ccr4 normalised against histone H3 (divided by cell volume (see Figure S1)) are represented as concentrations. **C)** Quantification ratios from the westerns for Xrn1 and Ccr4 over Rpb3. At least four biological replicates were used per GR. Data represent median and standard deviation. A Multiple Regression model was used for the statistical analysis and the associated p-values are shown (statistical significance, p-value < 0.1).

These data indicate a higher proportion of cytoplasmic concentrations of Xrn1 and Ccr4 compared to their association with chromatin, but the chromatin-associated forms increased less than RNA pol II. Therefore at a higher GR, the action of these factors seems to be more intensively connected to cytoplasmic mRNA degradation than to RNA pol II-dependent transcription. This finding agrees with the higher mRNA decay rates detected in the faster growing cells [4].

### TFIIS association with elongating RNA pol II increases with the growth rate

A relative decrease in RNA pol II-associated Xrn1 and Ccr4 provokes transcription elongation alterations characterised by higher RNA pol II backtracking frequency [25, 26]. We wondered whether their relative lower association at a higher GR might cause higher RNA pol II backtracking levels under these conditions. To corroborate the hypothesis of increased backtracking of elongating RNA pol II with the GR, we examined the association of TFIIS with chromatin in the same samples. It is well-known that TFIIS is required for the efficient reactivation of backtracked RNA pol II [27], and its recruitment to chromatin is a useful proxy of RNA pol II backtracking [26]. As depicted in Figure 7A and B, the concentration of the chromatin-associated TFIIS very significantly increased with the GR, but not in the WCEs. Notably, the chromatin-associated TFIIS to Ser2P-phosphorylated RNA pol II ratio was higher than it was vs. total Rpb3 (Figure 7C). This agrees with the lower proportion of the Ser2-phosphorylated enzymes in the chromatin of the high GR samples. These data support the notion that the alteration of RNA pol II elongation with the GR is likely accompanied by increased backtracking.

**Figure 7.**
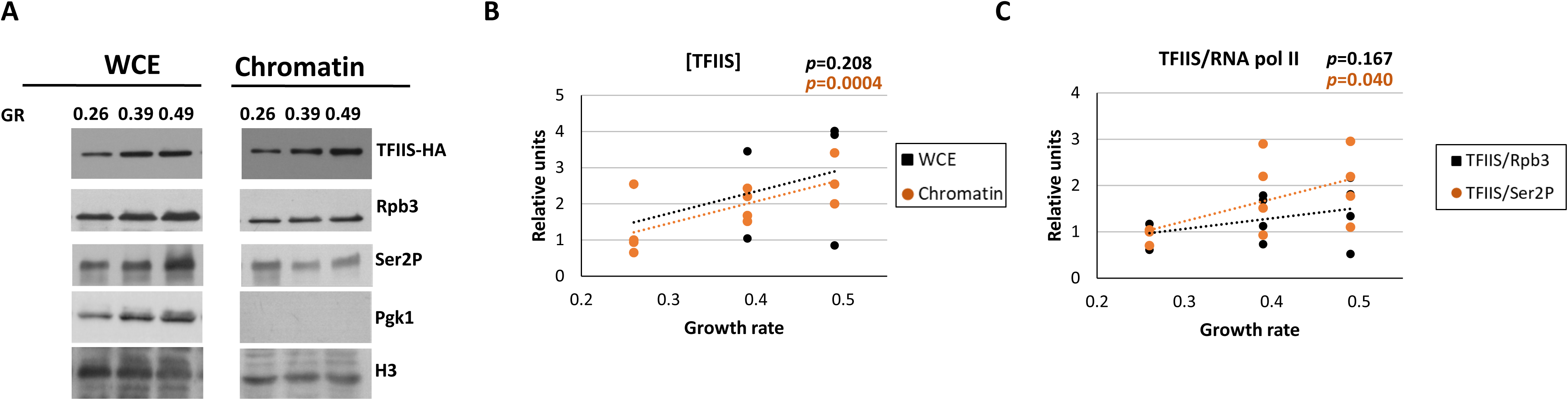
Elongation transcription factor TFIIS levels in chemostat cultures at different growth rates. **A)** The TFIIS (TFIIS-HA) levels from chemostat cultures at different growth rates (GRs) were determined by western blot analysis in the chromatin-enriched fractions and whole-cell crude extracts (WCEs) isolated by the yChEFs procedure [14, 15]. Histone H3 and Pgk1 as internal controls of chromatin and cytoplasm, respectively. **B)** Quantification of the western blot analysis of TFIIS normalised against histone H3 (divided by cell volume (see Figure S1)) are represented as concentrations. **C)** Quantification ratios from the westerns for TFIIS/Rpb3 and TFIIS/Ser2P. At least four biological replicates were used per GR. Data represent median and standard deviation. A Multiple Regression model was used for the statistical analysis and the associated p-values are shown (statistical significance, p-value < 0.1).

To confirm this conclusion, we investigated the occupancy of TFIIS for individual genes using the samples from the same cultures as in the previously presented data. According to Figure 8, the TFIIS to Ser2-phosphorylated RNA pol II ratio in chromatin increased in all the analysed genes.

**Figure 8.**
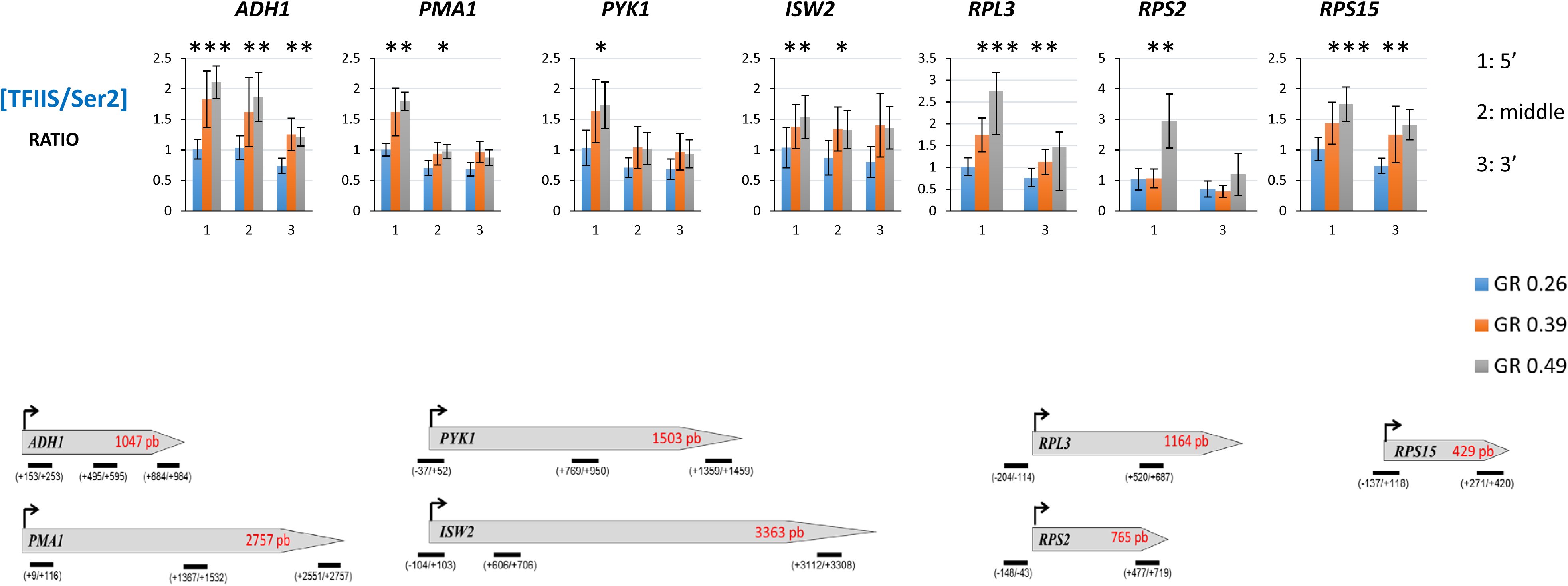
RNA pol II and RNA pol II CTD-Ser2phosphorylation levels in chemostat cultures at different growth rates, analysed by Chromatin Immunoprecipitation. Representation of the ratios between the data for TFIIS and Ser2P-CTD ChIPs analyses in different transcription units from the chemostat cultures at different growth rates (GRs). Data denote IP/INPUT ratios. At least three biological replicates were per GR. A Multiple Regression model was used for the statistical analysis and the associated p-values are shown (*: p-value < 0.1; **: p-value < 0.05; ***: p-value < 0.005).

## Discussion

Because it seems that cellular protein synthesis machinery (ribosomes, tRNAs and other factors) usually works at its maximum capacity (revised in [2] and other citations herein), any variation in the GR is accompanied by an increased amount of translation machinery [28, 29]. Given that RNA pol I and III are specifically devoted to transcribe rRNA and tRNAs, it is well-known that their activity (SRs) increases linearly with the GR [29]. In our experiments run with chemostat continuous cultures, we found that their association with chromatin increases with the GR (Figure 3). RNA pol II, however, transcribes thousands of different genes, most of which are not related to translation. In yeast, ribosomal proteins and other translation factors represent only about 15% of actual RNA pol II transcribing capacity [2, 30]. In fact the main RNA pol II products, mRNAs, do not need to be transcribed with an SR linked with the GR because they are unstable molecules that serve as templates for protein synthesis. Their influence on translation is related to their cellular concentrations, which depend on not only their SR, but also on their decay rates [31]. Variation in mRNA concentrations with GR in yeast has been extensively studied in the past, especially in D. Botstein’s lab [5–7, 10, 11]. Many mRNAs change their cellular concentrations with the GR. Thus it is especially relevant that these encoding ribosomal proteins and translation factors increase their concentrations with the GR [6, 7]. Most of those studies were done using chemostat cultures [10–12], where GRs can be tightly controlled and there is no change in growth conditions for long time periods. In these studies, however, it is not known if variation in mRNA levels is obtained by changes in the SR and/or in mRNA stabilities. In a previous study, in batch cultures we discovered that yeast cells increase the global RNA pol II SR with the GR, but maintain total [mRNA] constant because the global mRNA decay rate lowers [4]. In the present study, we analysed RNA pol II SR dependence with the GR under chemostat conditions to verify our previous conclusion. We found that the global amount of chromatin-associated and total cellular RNA pol II increased with the GR. (Figure 2). As RNA pol II is considered to have strong binding affinity to its gene targets, and that it is limiting for the transcription rate (see the discussion in refs. [22] and [32]), our results indicate that RNA pol II transcription regulation by the GR depends mostly on the control of the RNA pol II concentration, similarly as we found for its regulation by cell volume [22]. Interestingly, as dependence on the GR provokes an increase in RNA pol II levels and GR leads to, on the other hand, a smaller cell volume (Figure S1), the conclusion from these data would be that the SR increases with a decreasing cell volume. This is the opposite to what we observed for the relation between the SR and cell volume under other experimental conditions [23]. Nevertheless, this allows us to conclude that the rise in the SR due to the GR outweighs the expected decrease due to cell volume.

Our data also indicate that the GR alters the molecular properties of elongating RNA pol II by exhibiting lower Ser2- and Ser7-phosphorylation levels in its CTD domain (Figure 4C-D and 5). Ser2- and Ser7-phosphorylation are marks that influence RNA cleavage and polyadenylation, and also transcription termination [19]. Therefore, the altered form of elongating RNA pol II detected at a high GR might favour alternative polyA site selection, which has been previously connected with posttranscriptional effects, including altered mRNA stability [33]. Our previous work revealed that a high GR not only involves a rise in the SR by RNA pol II, but also increased mRNA turnover due to a parallel increase in mRNA decay [4].

Coordination between transcription and mRNA decay to control mRNA turnover without disrupting mRNA homeostasis is possible by the dual action of factors that can act in both transcription and mRNA decay [2, 8]. These factors include Xrn1 and Ccr4-Not, which have been demonstrated to independently stimulate RNA pol II elongation in addition to their previously known roles in mRNA degradation [25, 26, 34–36]. Our results indicated increased overall Xrn1 and Ccr4 concentrations with the GR, like that of chromatin-associated Ccr4. This agrees with the greater cytoplasmic mRNA decay demanded in high GRs [4].

A decrease in chromatin-associated Xrn1 involves higher RNA pol II backtracking frequency [25, 26, 35, 36]. Therefore at a high GR, it is conceivable that the nuclear scarcity of this factor compared to increased elongating RNA pol II levels would provoke a higher backtracking rate. In agreement with this prediction, we found greater recruitment of reactivating RNA cleavage factor TFIIS to elongating RNA pol II molecules with the GR (Figure 7C and 8).

In summary, our results confirm the overall response of RNA pol II-dependent transcription in yeast cells to the GR, and they indicate that this regulation involves molecular alterations of elongating RNA pol II molecules in CTD phosphorylation and backtracking.

## Materials and Methods

### Yeast strains and culture conditions

Yeast strain LMY5.2 (*MATa*, *TRP1::pADH::3HA::PPR2 trp1::kanMx4, his3*Δ*1, leu2*Δ*0, met15*Δ*0, ura3*Δ*0*) derived from BY4741 was used [13].

Chemostat cultures were established in a DASGIP parallel fermentation system (DASGIP AG, Jülich, Germany) equipped with four SR0400SS vessels. The working volume was set at 180 mL and YPD (2% glucose, 2% peptone, 1% yeast extract) was used as the growth medium. In all cases, agitation was maintained at 250 rpm and the temperature was set at 30°C using a water bath. Aerobic conditions were maintained by gassing the headspace of bioreactors with air (1.5 L/h). Off-gas was channelled through cooled condensers (1-3 °C) and the CO_2_ concentration in the exhaust gas was recorded every 30 s with a GA4 gas analyzer (DASGIP AG). Cultures were inoculated at 0.2 initial OD_600_ and continuous flow was started 9 h after inoculation. The out-pump was operated at a slightly higher speed of the in-pump so that the outlet pipe position would determine the vessel’s working volume. Peristaltic pumps were calibrated before and verified after each experiment, while the actual working volume was confirmed by the end of each experiment. These data were used to recalculate the actual dilution rates. Steady states were sampled only after continuous cultures had been running for at least six residence times and the CO_2_ production rate was constant. Samples were centrifuged at 4,500 g for 5 min. Cells and supernatants were frozen separately at −80°C and −20°C, respectively.

The glucose and ethanol contents in supernatants were quantified by a Surveyor Plus liquid chromatograph (Thermo Fisher Scientific, Waltham, MA) equipped with a refraction index (Surveyor RI Plus and Surveyor) on a 300 × 7.7 mm PL Hi-Plex H+ (8 μm particle size) column (Agilent Technologies, Santa Clara, CA, USA) and 4 × 3 mm ID Carbo-H guard (Phenomenex, Torrance, CA, USA). The column was maintained at 50°C and 1.5 mM H_2_SO_4_ was used as the mobile phase at a flow rate of 0.6 mL/min. Before injection in duplicate, samples were filtered through 0.22 μm pore size nylon filters.

### Flow cytometry

Frozen cells corresponding to 1 ml of cultures grown in chemostat at different GRs were used to analyse cell volume by flow cytometry. Cell pellets were fixed with 1 ml of absolute ethanol overnight, centrifuged and resuspended in 1 ml of 50 mM Tris-HCl pH 7.5. Cells were sonicated in a Bioruptor sonicator (Diagenode) for 3 cycles (30 s on / 30 s off cycles) for complete solubilisation and analysed by flow cytometry with an LSR Fortessa cell analyzer (BD). The results were explored using the FACSDiva software. The median of FSC-A was employed for representing the relative cell volume between different conditions.

### Chromatin-enriched fractions and western blot analyses

Chromatin-enriched fraction purification was performed according to the yChEFs procedure [14, 15] using 36 OD units of frozen cells from cultures grown in a chemostat at different GRs. The final chromatin-bound proteins were resuspended in 1X SDS-PAGE sample buffer, boiled and analysed by western blot with different antibodies.

Protein electrophoresis and western blots were carried out as described in [37]. The employed antibodies included anti-Rpb1 (against the CTD; manufactured in our laboratory) [14], anti-Rpb1 (y-80; Santa Cruz Biotechnology), anti-RNA pol II phospho Ser7 (4E12; Chromotek), anti-RNA pol II phospho Ser5 (CTD4H8; Millipore), anti-RNA pol II phospho Ser2 (ab5095; Abcam), anti-Rpb3 (anti-POLR2C;1Y26, Abcam), anti-Rpb4 (Pol II RPB4 (2Y14); Biolegend), anti-haemagglutinin (anti-HA; 12CA5; Roche), anti-phosphoglycerate kinase, Pgk1 (22C5D8; Invitrogen), anti-H3 (ab1791; Abcam), anti-A34.5 and anti-C128 (gifts from P. Thuriaux), anti-Xrn1 (gift from Arlen Johnson) and anti-Ccr4 (gift from Martine Collart).

The IMAGE STUDIO LITE software was utilised to quantify the intensities of the immunoreactive bands.

### Chromatin immunoprecipitation

For the ChIP procedure, live cells must be treated with formaldehyde to provoke protein-protein and protein-DNA cross-links between molecules, which are in close proximity on the chromatin template *in vivo* [38]. The samples from the chemostat cultures used in this work were frozen at −80°C without formaldehyde treatment. Then the frozen cells for each GR (from 0.26 to 0.49 h^-1^) were thawed, treated with 1% formaldehyde and later subjected to chromatin sonication. Strikingly, we noted that the amount of retrieved chromatin decreased with the GR (Figure S2A). We suspected that the physiological state of the yeast cells at a higher GR, followed by flash-freezing, provoked overcross-linking with formaldehyde, which would later decrease the amount of purified DNA. Therefore, we assayed a lower formaldehyde concentration (0.25%) during a trial experiment, in which cells were previously frozen for 40 days at −80°C. In that case, chromatin was properly recovered, which was not the case for 1% formaldehyde (Figure S2B). As 0.25% formaldehyde was enough to also achieve good cross-linking at low GRs, we performed ChIP experiments with the whole set of chemostat cultures for all the GRs after thawing cells and treating them with 0.25% formaldehyde to generate cross-linking.

Chromatin immunoprecipitations were performed as previously described [37], with some modifications. Briefly, 25 OD units of frozen cells from the cultures grown in a chemostat at different GRs were used. Cells were resuspended in 50 ml Tris-saline buffer (150 mM NaCl, 20 mM Tris-HCl, pH 7.5) and cross-linking was carried out by adding 0.4% formaldehyde (v/v) to cells and incubating them at room temperature for 15 min with some agitation. Then 2.5 ml of 2.5 M glycine were added and cells were incubated for 5 min at room temperature. Cells were harvested and washed 4 times with 25 ml Tris–saline buffer (150 mM NaCl, 20 mM Tris-HCl, pH 7.5) at 4°C. Cell breakage was performed in 300 µl lysis buffer (50 mM HEPES, 140 mM NaCl, 1 mM EDTA, pH 8, 1% (v/v) Triton-X-100, 0.1% (w/v) sodium deoxycholate, 1 mM PMSF, 0.15% (w/v) Benzamidine, protease inhibitor cocktail (Complete, Roche)) with glass beads (425-600 µm). Cell extracts were sonicated in a Bioruptor sonicator (Diagenode) for 30 min during 3 cycles of 10 min (30 s/on / 30 s off cycles) at the highest power (chromatin was sheared into an average size of 300 bp). Extracts were clarified at 12,000 rpm for 5 minutes at 4°C. Finally, 50 µl of chromatin were used for each chromatin immunoprecipitation experiment, while 10 µl were utilised for the input samples.

Rpb3-chromatin immunoprecipitation was conducted using Dyna-beads M-280 sheep anti-mouse IgG (BioRad) and anti-Rpb3 (anti-POLR2C;1Y26, Abcam). Ser2-chromatin immunoprecipitation was performed employing Dyna-beads M-280 sheep anti-rabbit IgG (BioRad) and anti-RNA pol II phospho Ser2 (ab5095; Abcam). TFIIS-HA-chromatin immunoprecipitation was conducted with Dyna-beads sheep anti-rat IgG (BioRad) and anti-HA high affinity (3F10; Roche).

Real-time PCR was run in a CFX-384 Real-Time PCR instrument (BioRad) with SYBR premix EX Taq (Takara) and the oligonucleotides listed in Table 1. At least three independent biological replicates were analysed. For quantitative real-time PCR, 1:100 dilution was applied for the input DNA.

**Table 1.**
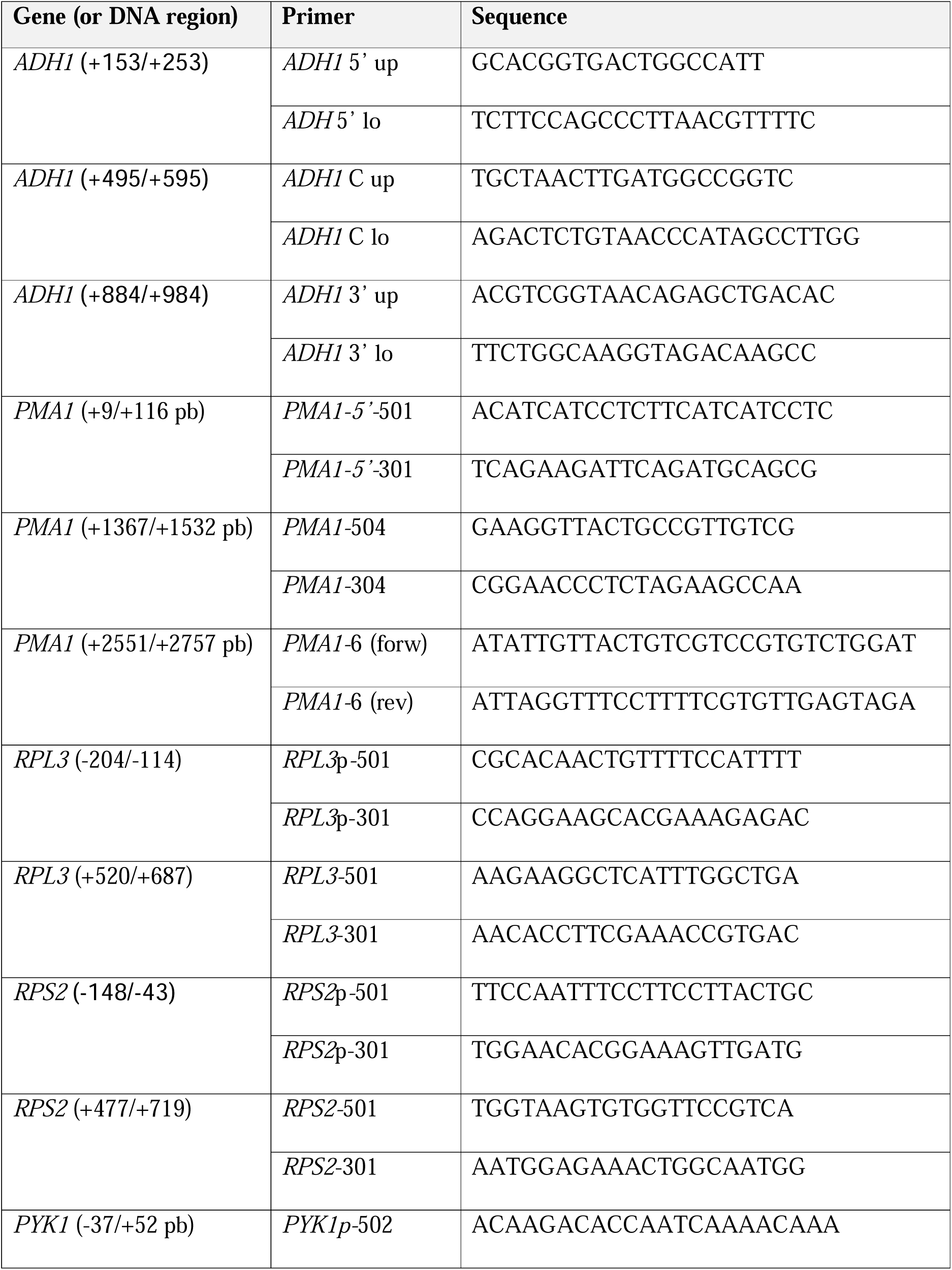
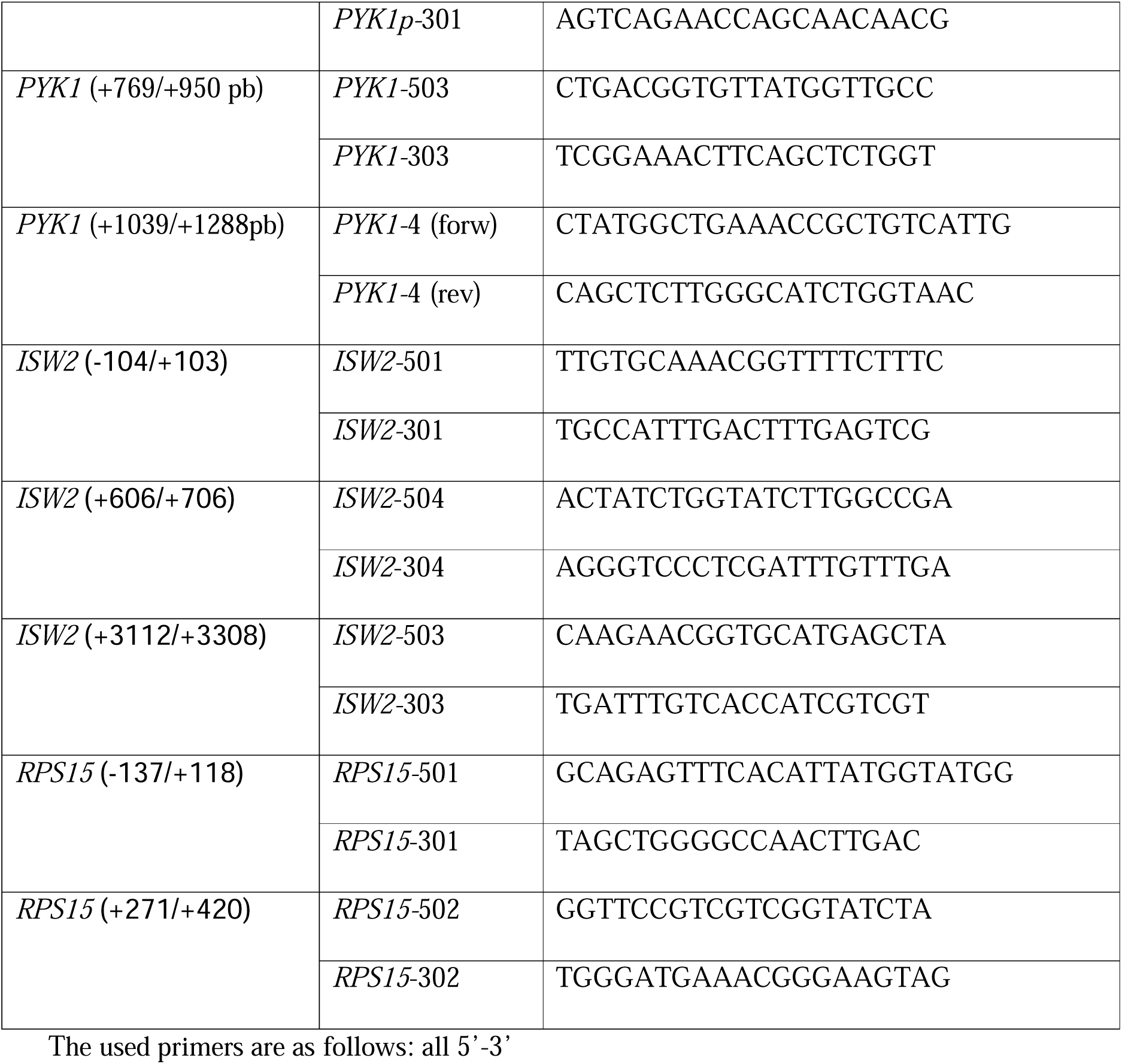
Oligonucleotides used.

## Funding

This work has been supported by grants from the Spanish Ministry of Science and Innovation (MCIN) and ERDF to F.N. (PID2020-112853GBC33), J.E.P-O (PID2020-112853GBC31) and S.Ch. (PID2020-112853GB-C32), by the Spanish Ministry of Science, Innovation and Universities (MCIU) and ERDF to F.N. (PID2023-148037NB-C22) and S.Ch. (PID2023-148037NB-C21), by the Junta de Andalucía (BIO258) to F.N. and S.Ch (BIO271) and by the Spanish Ministry of Science and Innovation (MICINN) and ERDF to F.N., J.E.P-O and S. Ch. (RED2018-102467-T).

## Acknowledgement

We thank the ‘Servicios Centrales de Apoyo a la Investigación (SCAI)’ of the Universidad de Jaén for technical support.

## Supplementary Figure Legends

**Figure S1. Cell size determination.** The cells from the chemostat cultures at different growth rates (GRs) were subjected to a flow cytometry analysis and cell size was determined by the FSC-A median (left graph). The results were analysed with the FACSDiva software. At least four biological triplicates were used per GR.

**Figure S2. Analysis of sonicated chromatin. A)** Sonicated chromatin for the frozen cells from chemostat cultures at three different growth rates (GRs) with cross-linking using the 1% formaldehyde concentration. Three biological replicates for each analysis are shown. **B)** Sonicated chromatin in frozen and unfrozen cells with cross-linking at different formaldehyde concentrations. Two biological replicates for each analysis are shown.

## Notes

### Competing Interest Statement

The authors have declared no competing interest.

## Bibliography

[1] D.W. Thompson, On Growth and Form, in, Cambridge University Press, Cambridge, 1917.

[2] J.E. Pérez-Ortín, V. Tordera, S. Chávez, Homeostasis in the Central Dogma of Molecular Biology: the importance of mRNA instability, RNA Biol, 16 (2019) 1659–1666.

[3] G.M. Caballero-Córdoba, V.C. Sgarbieri, Nutritional and toxicological evaluation of yeast (*Saccharomyces cerevisiae*) biomass and a yeast protein concentrate, J Sci Food Agr, 80 (2000) 341–351.

[4] J. Garcia-Martinez, L. Delgado-Ramos, G. Ayala, V. Pelechano, D.A. Medina, F. Carrasco, R. Gonzalez, E. Andres-Leon, L. Steinmetz, J. Warringer, S. Chavez, J.E. Perez-Ortin, The cellular growth rate controls overall mRNA turnover, and modulates either transcription or degradation rates of particular gene regulons, Nucleic Acids Res, 44 (2016) 3643–3658.

[5] N. Slavov, D. Botstein, Coupling among growth rate response, metabolic cycle, and cell division cycle in yeast, Molecular Biology of the Cell, 22 (2011) 1997–2009.

[6] M.J. Brauer, C. Huttenhower, E.M. Airoldi, R. Rosenstein, J.C. Matese, D. Gresham, V.M. Boer, O.G. Troyanskaya, D. Botstein, Coordination of growth rate, cell cycle, stress response, and metabolic activity in yeast, Mol Biol Cell, 19 (2008) 352–367.

[7] E.M. Airoldi, C. Huttenhower, D. Gresham, C. Lu, A.A. Caudy, M.J. Dunham, J.R. Broach, D. Botstein, O.G. Troyanskaya, Predicting cellular growth from gene expression signatures, PLoS Comput Biol, 5 (2009) e1000257.

[8] S. Chávez, J. García-Martínez, L. Delgado-Ramos, J.E. Pérez-Ortín, The importance of controlling mRNA turnover during cell proliferation, Curr Genet, 62 (2016) 701–710.

[9] J. García-Martínez, K. Troule, S. Chávez, J.E. Pérez-Ortín, Growth rate controls mRNA turnover in steady and non-steady states, RNA Biol, 13 (2016) 1175–1181.

[10] M.J. Brauer, A.J. Saldanha, K. Dolinski, D. Botstein, Homeostatic adjustment and metabolic remodeling in glucose-limited yeast cultures, Molecular biology of the cell, 16 (2005) 2503–2517.

[11] A.J. Saldanha, D. Brauer Mj Fau - Botstein, D. Botstein, Nutritional homeostasis in batch and steady-state culture of yeast, in: Mol Biol Cell., vol. 15, pp. 4089–4104. doi: 4010.1091/mbc.e4004-4004-0306. Epub 2004 Jul 4087.

[12] M.J. Dunham, E.O. Kerr, A.W. Miller, C. Payen, Chemostat Culture for Yeast Physiology and Experimental Evolution, Cold Spring Harb Protoc., 2017 pdb.top077610. doi: 077610.071101/pdb.top077610.

[13] F. Gómez-Herreros, L. de Miguel-Jiménez, M. Morillo-Huesca, L. Delgado-Ramos, M.C. Muñoz-Centeno, S. Chávez, TFIIS is required for the balanced expression of the genes encoding ribosomal components under transcriptional stress, Nucleic Acids Res, 40 (2012) 6508–6519.

[14] A. Cuevas-Bermúdez, A. Garrido-Godino, F. Navarro, A novel yeast chromatin-enriched fractions purification approach, yChEFs, for the chromatin-associated protein analysis used for chromatin-associated and RNA-dependent chromatin-associated proteome studies from *Saccharomyces cerevisiae*, Gene Reports, 16 (2019) 100450.

[15] A. Cuevas-Bermúdez, A.I. Garrido-Godino, F. Gutiérrez-Santiago, F. Navarro, A Yeast Chromatin-enriched Fractions Purification Approach, yChEFs, from *Saccharomyces cerevisiae*, Bio-protocol, 10 (2020) e3471.

[16] K.L. Claude, D. Bureik, D. Chatzitheodoridou, P. Adarska, A. Singh, K.M. Schmoller, Transcription coordinates histone amounts and genome content, Nat Commun, 12 (2021) 4202.

[17] J.E. Pérez-Ortín, D.A. Medina, S. Chávez, J. Moreno, What do you mean by transcription rate?: the conceptual difference between nascent transcription rate and mRNA synthesis rate is essential for the proper understanding of transcriptomic analyses, Bioessays, 35 (2013) 1056–1062.

[18] W.H. Mager, R.J. Planta, Coordinate expression of ribosomal protein genes in yeast as a function of cellular growth rate, Mol Cell Biochem, 104 (1991) 181–187.

[19] H. Kim, B. Erickson, W. Luo, D. Seward, J.H. Graber, D.D. Pollock, P.C. Megee, D.L. Bentley, Gene-specific RNA polymerase II phosphorylation and the CTD code, Nat Struct Mol Biol, (2010).

[20] M. Kim, H. Suh, E.J. Cho, S. Buratowski, Phosphorylation of the yeast Rpb1 C-terminal domain at serines 2, 5, and 7, J Biol Chem, 284 (2009) 26421-26426.

[21] P. Allepuz-Fuster, V. Martínez-Fernández, A.I. Garrido-Godino, S. Alonso-Aguado, S.D. Hanes, F. Navarro, O. Calvo, Rpb4/7 facilitates RNA polymerase II CTD dephosphorylation, Nucleic Acids Res, 42 (2014) 13674–13688.

[22] A. Mena, D.A. Medina, J. García-Martínez, V. Begley, A. Singh, S. Chávez, M.C. Muñoz-Centeno, J.E. Pérez-Ortín, Asymmetric cell division requires specific mechanisms for adjusting global transcription, Nucleic Acids Res, 45 (2017) 12401–12412.

[23] M.A. Collart, The Ccr4-Not complex is a key regulator of eukaryotic gene expression, Wiley Interdiscip Rev RNA, 7 (2016) 438–454.

[24] D.A. Medina, A. Jordán-Pla, G. Millán-Zambrano, S. Chávez, M. Choder, J.E. Pérez-Ortín, Cytoplasmic 5’-3’ exonuclease Xrn1p is also a genome-wide transcription factor in yeast, Front Genet, 5 (2014) 1.

[25] V. Begley, A. Jordán-Pla, X. Penate, A.I. Garrido-Godino, D. Challal, A. Cuevas-Bermudez, A. Mitjavila, M. Barucco, G. Gutiérrez, A. Singh, P. Alepuz, F. Navarro, D. Libri, J.E. Pérez-Ortín, S. Chávez, Xrn1 influence on gene transcription results from the combination of general effects on elongating RNA pol II and gene-specific chromatin configuration, RNA Biol, (2020) 1–14.

[26] V. Begley, D. Corzo, A. Jordán-Pla, A. Cuevas-Bermúdez, L. Miguel-Jiménez, D. Pérez-Aguado, M. Machuca-Ostos, F. Navarro, M.J. Chávez, J.E. Pérez-Ortín, S. Chávez, The mRNA degradation factor Xrn1 regulates transcription elongation in parallel to Ccr4, Nucleic Acids Res, 47 (2019) 9524–9541.

[27] F. Gómez-Herreros, L. de Miguel-Jiménez, G. Millán-Zambrano, X. Peñate, L. Delgado-Ramos, M.C. Muñoz-Centeno, S. Chávez, One step back before moving forward: regulation of transcription elongation by arrest and backtracking, FEBS Lett, 586 (2012) 2820–2825.

[28] J.R. Warner, The economics of ribosome byosynthesis in yeast, Trends Biochem Sci 24 (1999) 437–440.

[29] E. Metzl-Raz, M. Kafri, G. Yaakov, N. Barkai, Gene Transcription as a Limiting Factor in Protein Production and Cell Growth, G3 Genes|Genomes|Genetics, 10 (2020) 3229-3242.

[30] V. Pelechano, S. Jimeno-González, A. Rodríguez-Gil, J. García-Martínez, J.E. Pérez-Ortín, S. Chávez, Regulon-specific control of transcription elongation across the yeast genome, PLoS Genet, 5 (2009) e1000614.

[31] J.E. Pérez-Ortín, P.M. Alepuz, J. Moreno, Genomics and gene transcription kinetics in yeast, Trends Genet, 23 (2007) 250–257.

[32] J.E. Pérez-Ortín, M.J. García-Marcelo, I. Delgado-Román, M.C. Munoz-Centeno, S. Chávez, Influence of cell volume on the gene transcription rate, Biochim Biophys Acta Gene Regul Mech, 1867 (2024) 195008.

[33] I. Gupta, S. Clauder-Munster, B. Klaus, A.I. Jarvelin, R.S. Aiyar, V. Benes, S. Wilkening, W. Huber, V. Pelechano, L.M. Steinmetz, Alternative polyadenylation diversifies post-transcriptional regulation by selective RNA-protein interactions, Mol Syst Biol, 10 (2014) 719.

[34] J. Fischer, Y.S. Song, N. Yosef, J. di Iulio, L.S. Churchman, M. Choder, The yeast exoribonuclease Xrn1 and associated factors modulate RNA polymerase II processivity in 5’ and 3’ gene regions, The Journal of biological chemistry, 295 (2020) 11435–11454.

[35] A. Dutta, V. Babbarwal, J. Fu, D. Brunke-Reese, D.M. Libert, J. Willis, J.C. Reese, Ccr4-Not and TFIIS Function Cooperatively To Rescue Arrested RNA Polymerase II, Mol Cell Biol, 35 (2015) 1915–1925.

[36] J.A. Kruk, A. Dutta, J. Fu, D.S. Gilmour, J.C. Reese, The multifunctional Ccr4-Not complex directly promotes transcription elongation, Genes Dev, 25 (2011) 581–593.

[37] A.I. Garrido-Godino, M.C. García-López, F. Navarro, Correct assembly of RNA polymerase II Depends on the foot domain and Is required for multiple steps of transcription in *Saccharomyces cerevisiae*, Mol Cell Biol, 33 (2013) 3611–3626.

[38] O. Aparicio, J.V. Geisberg, K. Struhl, Chromatin immunoprecipitation for determining the association of proteins with specific genomic sequences in vivo, in: Curr Protoc Cell Biol, vol. Chapter 17, 2004, pp. Unit 17 17.

