## Supplementary figures and images for "Growth rate regulates RNA polymerase II transcription elongation in *Saccharomyces cerevisiae*"

### Supplemental Figures

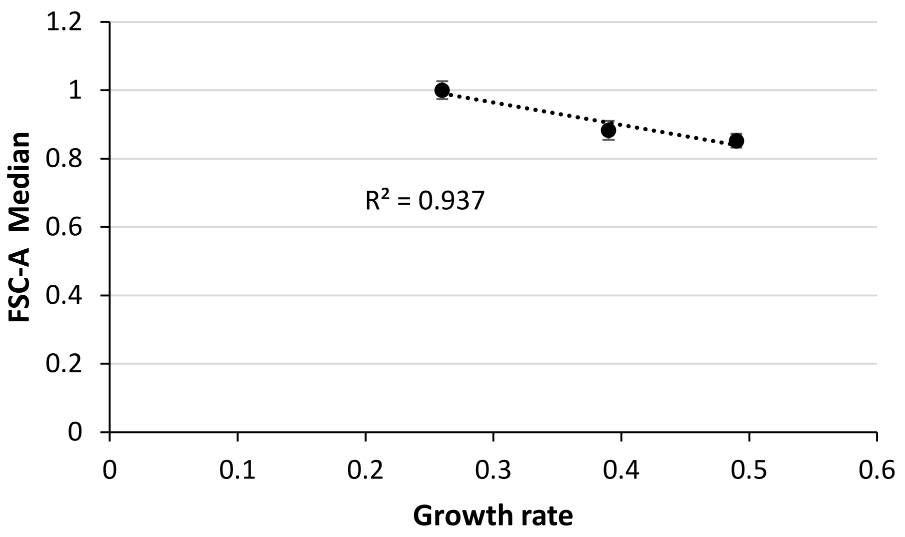

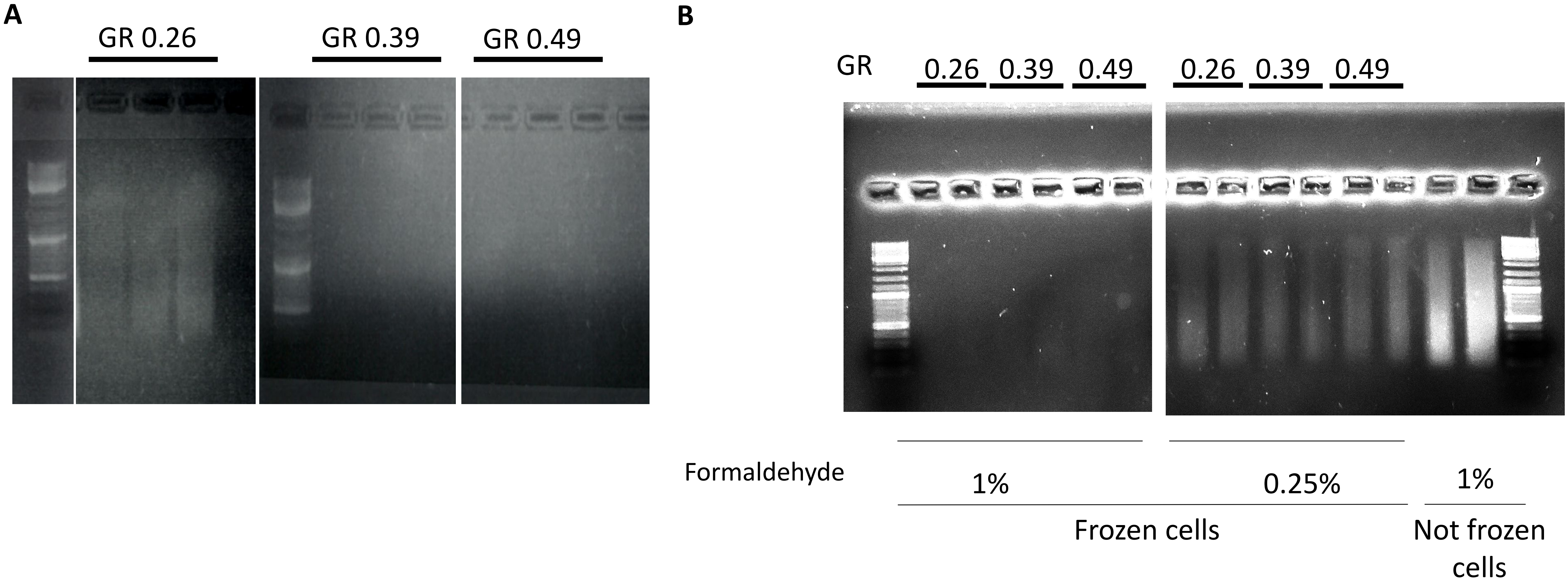
